# Metagenomic ecosystem monitoring of soft scale and mealybug infestations in Australian vineyards

**DOI:** 10.1101/2023.07.30.551191

**Authors:** Chris M. Ward, Cristobal A. Onetto, Steven Van Den Heuvel, Robyn Dixon, Anthony R. Borneman

## Abstract

Soft scale insects and mealybugs are phloem feeding Hemipterans that are considered majors pests in agricultural and horticultural settings throughout the world. Viticulturally, scale are a major issue due to their ability to secrete honeydew, which facilitates the development of sooty mould and for their propensity as transmission vectors for several viral diseases of grapevine. To facilitate the rapid identification and quantification of vineyard-associated insects a metagenomic-based bioinformatic pipeline was developed for generalised ecosystem monitoring that automated the assembly and classification of insect mitochondrial genomes from shotgun sequencing data using the Barcode of Life Database API. *Parthenolecanium corni* (European fruit scale), which was thought to be absent from Australian grapevines, was identified as the dominant coccid species infesting all vines sampled, along with secondary infestation by *Pseudococcus viburni* (obscure mealybug) and *Pseudo. longispinus* (long-tailed mealybug). In addition, parisitoidism by *Coccophagus scutellaris* (Aphelinidae) wasps was also detected. The discovery of *Parth. corni* as a significant member of scale infestations in Australia has significant implications for the development of effective control strategies for this important group of pests.

## Introduction

Worldwide, the family Coccidae (Coccoidea) contains greater than 1100 species classified into 163 genera, although only 80 species have been described as occurring within Australia (García Morales *et al*., 2016). Likewise, of the 2143 described Pseudococcidae (Coccoidea), only 210 have been detected as contemporary Australian populations (García Morales *et al*., 2016; Moir, 2021).

Several coccids have been shown to affect the grapevine *Vitis vinifera*, including various species within the genera *Ceroplastes, Coccus, Cryptinglisia, Eulecanium, Mesolecanium, Neopulvinaria, Neolecanium, Parasaissetia, Parthenolecanium, Pseudokermes, Pulvinaria, Saissetia* and *Trijuba* (Ben-Dov, 1977; Ben-Dov & Hodgson, 1997; Rakimov *et al*., 2013; Rakimov *et al*., 2015). Among these, *Parth. persicae* (European peach scale), *Parth. pruinosum* (frosted scale), and *Coccus hesperidum* (brown soft scale) are frequently encountered in Australian vineyards, while *Co. longulus* (long brown scale), *Parasaissetia nigra* (black scale), and *Saissetia* species have also been observed with lower frequency (Rakimov *et al*., 2013).

Recent surveys conducted in Australia to assess disease pressure and the distribution of Coccids indicate that scale insects are rapidly becoming a significant pest of grapevines (Essling, 2018; Venus, 2017). The distribution of scale and mealybugs appears to be uneven across different grape-growing regions, with Western Australia and Queensland showing less frequent infestations than other states (Rakimov *et al*., 2013).

While severe infestations of scale and mealybugs can have direct detrimental effects on vine growth through the depletion of nutrients, the main negative impacts of infestation are manifest through secondary effects. Firstly, the secretion of sugar-rich honeydew as a by-product of scale and mealybug feeding, facilitates the growth of sooty mould throughout the vine, which can lead to quality reductions and/or total loss of the grape crop (Ben-Dov & Hodgson, 1997).

In addition to promoting sooty mould, scale and mealybugs also represent known or suspected vectors of grapevine ampelo- and viti-viruses, including grapevine virus A (GVA) (Hommay *et al*., 2008) and grapevine-leafroll associated viruses (GLRaV) 1 and 3 (Bahder *et al*., 2013; Sforza *et al*., 2003). Infection by these viruses can have devastating effects on yield and fruit quality, including reduced accumulation of sugars, increased acidity and irregular flavour profiles (Goheen & Cook, 1959; Naidu *et al*., 2014). The mobile nymph stages of both types of insects are suspected to transmit these viruses between adjacent vines, although recent studies indicate that passive aerial dispersal of the nymphs can promote long-distance dispersal across a vineyard (Barrass *et al*., 1994; Hommay *et al*., 2019).

Studies have demonstrated the existence of several natural enemies of coccids in Australia, including parasitoid wasps, beetles, moth larvae, and lacewings (Rakimov *et al*., 2013; Rakimov *et al*., 2015). Therefore, chemical control strategies for coccids are only recommended in severe infestation cases to avoid harming the beneficial parasitoids that inhabit the vineyards (Frank, 2012; Rakimov *et al*., 2015).

Ecosystem monitoring utilizing molecular or next generation sequencing methodologies have been used to successfully characterize beetle species richness (Crampton-Platt *et al*., 2015), airborne fungal diversity (Banchi *et al*., 2018), and track pollinator populations (Harper *et al*., 2023). However, successful species classification generally requires strong genetic databases to first be developed to carry out high-throughput monitoring (Crampton-Platt *et al*., 2015). Targeted ecosystem monitoring was carried out previously by Rakimov *et al*. (2013), which utilized a COX1 amplicon to identify individual soft scale insects that infested grapevines across several Australian vineyards. The results of this work concluded that *Parth. persicae* was the dominant coccid present, with secondary infestations commonly observed by *Parth. pruinosum*.

In order to facilitate the rapid identification and quantification of vineyard-associated insects an untargeted ecosystem monitoring protocol was developed and its utility demonstrated by determining the composition of insect populations associated with scale infestation. Shotgun metagenomic sequencing was employed against population scrapings from areas of soft scale-infestation from vines across four wine regions within South Australia. To enable the high-throughput analysis of the large datasets produced by this approach, a rapid pipeline to assemble and classify insect mitochondrial genomes present was established that takes advantage of the Barcode of Life Database (BOLD) API (Ratnasingham & Hebert, 2007).

## Materials and methods

### 1. Collection and DNA extraction and sequencing

Samples were collected from eight vineyards across South Australia (Table 1) by taking scrapings at infestation sites from scale-infested vines. Scrapings were immediately stored under ethanol to preserve tissue. DNA isolation was carried out by grinding solid material using a Geno/Grinder 2010 (SPEX), followed by extraction using the DNEasy Plant Kit (Qiagen).

**Table 1.** Assembly statistics for each metagenomic sample.

| Sample | Location | Contigs | Total<br>length<br>(kb) | Longest<br>contig | N<sub>50</sub> | Full<br>mitochondria<br>l genomes | Unique<br>insect<br>taxa |
| --- | --- | --- | --- | --- | --- | --- | --- |
| 7361 | Langhorne Creek | 452 | 425.6 | 44283 | 955 | 4 | 6 |
| 7372 | McLaren Vale | 305 | 212.5 | 10753 | 747 | 0 | 3 |
| 7373 | McLaren Vale | 963 | 787.4 | 12427 | 1023 | 1 | 2 |
| 7374 | Langhorne Creek | 910 | 735.4 | 15563 | 1078 | 1 | 1 |
| 7375 | Adelaide Hills | 873 | 717.9 | 15198 | 1124 | 1 | 2 |
| 7376 | Adelaide Hills | 1015 | 786.6 | 15297 | 900 | 1 | 1 |
| 7377 | Barossa Valley | 940 | 689.8 | 6523 | 861 | 1 | 2 |
| 7378 | Barossa Valley | 981 | 763.3 | 15429 | 911 | 1 | 2 |

Sequencing libraries were prepared using the Illumina DNA library kit and sequenced on an Illumina NovaSeq 6000 using 2 × 150bp chemistry (Ramaciotti Centre for Genomics, University of New South Wales, Sydney, Australia). Quality control was carried out on short read data using fastqc (https://www.bioinformatics.babraham.ac.uk/projects/fastqc/) and ngsReports (Ward, To, *et al*., 2020).

All sequencing data can be obtained from NCBI under the BioProject accession PRJNA986325.

### 2. Genome assembly and analysis

Mitochondrial genome assembly was first carried out by mapping raw sequence reads to all available, complete insect mitochondrial genomes (n = 12880) present in NCBI GenBank using BWA-mem (Li, 2013) and sorted using SAMtools (Li *et al*., 2009); removing all unmapped reads. Read IDs of mitochondrial mapped reads were then used to extract putative mitochondrial reads from the original *fastq* files using seqtk subseq (Li, 2012). Putative mitochondrial reads were then assembled using megahit under default settings. Contigs were then remapped back to the RefSeq insect mitochondrial genome database using minimap2 (Li, 2018) with the ‘splice’ alignment preset. Contigs that were successfully mapped to the insect mitochondrial database were then extracted using seqkit subseq (Shen *et al*., 2016). Each mitochondrial contig was then annotated independently using MITOS (Bernt *et al*., 2013) using the insect mitochondrial genetic code and MITOS reference database.

COX1 sequences were extracted from each of the contig annotations and were provided as a seed sequence to NOVOplasty (Dierckxsens *et al*., 2017) to carry out circularised assembly of the mitochondrial genome using the raw short reads. Mitochondrial genes, rRNAs, and tRNAs were then annotated within each circularised genome using MITOS (Bernt *et al*., 2013) with the insect mitochondrial genetic code and MITOS reference database.

### 3. Species assignment and phylogenetics

Taxa identification was carried out using two methods. First, COX1 sequences were extracted from all putative mitochondrial contigs and queried against the ‘COX1’ BOLD database in a highly parallelised manner using the BOLD.R (Mudalige, 2021) and futures parallelisation (Bengtsson, 2020) R packages to pass sequences to the BOLD API (Ratnasingham & Hebert, 2007). Second, mitochondrial contigs were queried against the RefSeq Insect mitochondrial genome database from above using BLASTn (Camacho *et al*., 2009).

COX1 sequences identified as *Parthenolecanium corni* using the BOLD database were extracted using geaR (Ward, Ludington, *et al*., 2020) and aligned to the two COX1 amplicon loci available for this genus using MAFFT (Katoh & Standley, 2013) and maximum likelihood phylogenies were constructed using the best fit model argument in IQ-TREE (Nguyen *et al*., 2015).

To provide further evidence that *Parthenolecanium sp*. from the Langhorne Creek (7361) scraping were *Parth. corni*, individuals were visually assessed for taxonomic characters, including dorsal tubular duct density and tubular duct presence on the dorsum of the anal cleft (Gill, 1988), that have been stated to separate *Parth. pruinosum* and *Parth. corni*.

### 4. MitoMiner pipeline

The MitoMiner pipeline was written utilizing the snakemake workflow framework and can be accessed at github.com/AWRI/MitoMiner

## Results and Discussion

### Highly parallelised, automated assembly of insect mitochondrial genomes from metagenomic samples using MitoMiner

Ecosystem monitoring using high throughput genomic methods is a powerful tool to deconvolute the composition of mixed population samples (Banchi *et al*., 2018; Crampton-Platt *et al*., 2015; Harper *et al*., 2023). To apply this methodology to investigate the composition of insect populations associated with scale infestation, shotgun metagenomic sequencing was employed against population scrapings from areas of soft-scale infestation from vines across four wine regions within South Australia. Individual grapevines were surveyed for scale insect infestation across eight vineyards from across Langhorne Creek, Adelaide Hills, Barossa Valley and McLaren Vale (Table 1), with an average of 28.9 ± 14 Gb of sequencing data produced across the samples.

To facilitate the genome assembly, annotation and species assignment required to transform the shotgun sequencing data into species and abundance assignments, the MitoMiner pipeline was developed. MitoMiner is a snakemake implementation that extracts putative mitochondrial reads from short read data and assembles them utilizing a metagenomic assembly method in megahit. Putative mitochondrial contigs are then automatically annotated, circularised and classified to taxonomic origin utilizing information present in databases from both NCBI and BOLD (Figure 1).

**Figure 1.**
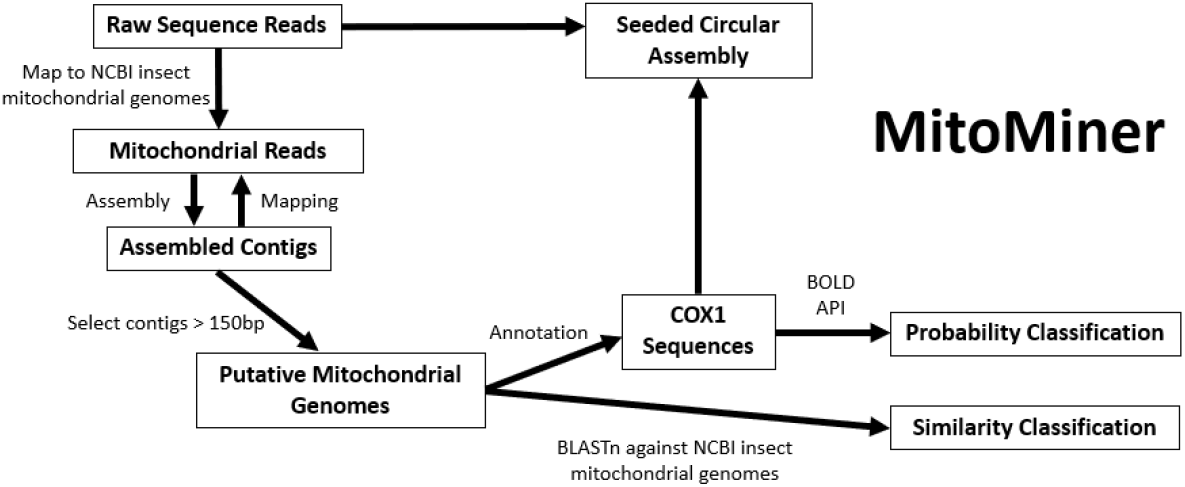
Workflow schematic of the MitoMiner pipeline implemented in snakemake.

After mapping raw reads against the NCBI insect mitochondrial genome database an average of 2.5 ± 0.78 M reads could be confidently classified as having a mitochondrial origin and were retained for subsequent assembly and analysis. Assembled putative mitochondrial contigs ranged in size from 0.22 to 44.2 kb and an average of 804 ± 270 mitochondrial contigs were recovered per metagenomic assembly (Table 1). Mitochondrial annotation identified a COX1 coding sequence present in 4.9 % of all mitochondrial contigs, suggesting many contigs are likely fragments derived from the same species (Table 1).

Classification of COX1 against BOLD identified seven Insect taxa across all samples, confidently (≥ 0.95 probability) assigning six to species level. COX1 sequences classified to any insect lineages were then utilized as seed sequences for circularized mitochondrial genome assembly, resulting in circularized complete assemblies for four unique taxa (Table 1). Species assignment of mitochondrial contigs was then performed to ascertain the species distribution and frequency across the vineyard samples.

### *Parthenolecanium* and *Pseudococcus* species co-occur in the same niche and are being actively targeted by parasitoids

Across the eight vineyard samples, complete mitochondrial genomes, containing 12 protein coding genes and 2 rRNA arrays, were recovered for *Parth. corni* (15.5 kb), *Pseudo. viburni* (15.2 kb), *Anatrachyntis badia* (Lepidoptera) (15.4 kb) and *Corticarina gibbosa* (Coleoptera) (16.1 kb). Whereas partial mitochondrial genomes were recovered for *Coccophagus scutellaris* (Hymenoptera) (6.5 kb), and *Pseudo. longispinius* (0.88 kb).

When considering individual samples, multiple Hemipteran pest species were identified within the same scraping, suggesting that soft scale and mealybugs can co-occur at close proximity (Figure 2). Only a single insect species, *Parth. corni*, was identified across all 8 locations (Figure 2), with all 8 samples sharing 100 % DNA identity across the mitochondrial COX1 locus. Three of the sites sampled were predicted to contain *Pseudococcus* species, with two samples being classified as containing *Pseudo. vibruni* (7361, Langhorne Creek; 7372, McLaren Vale), while the third was predicted to contain mitochondrial DNA from *Pseudo. longispinius* (7373, McLaren Vale) (Figure 2).

**Figure 2.**
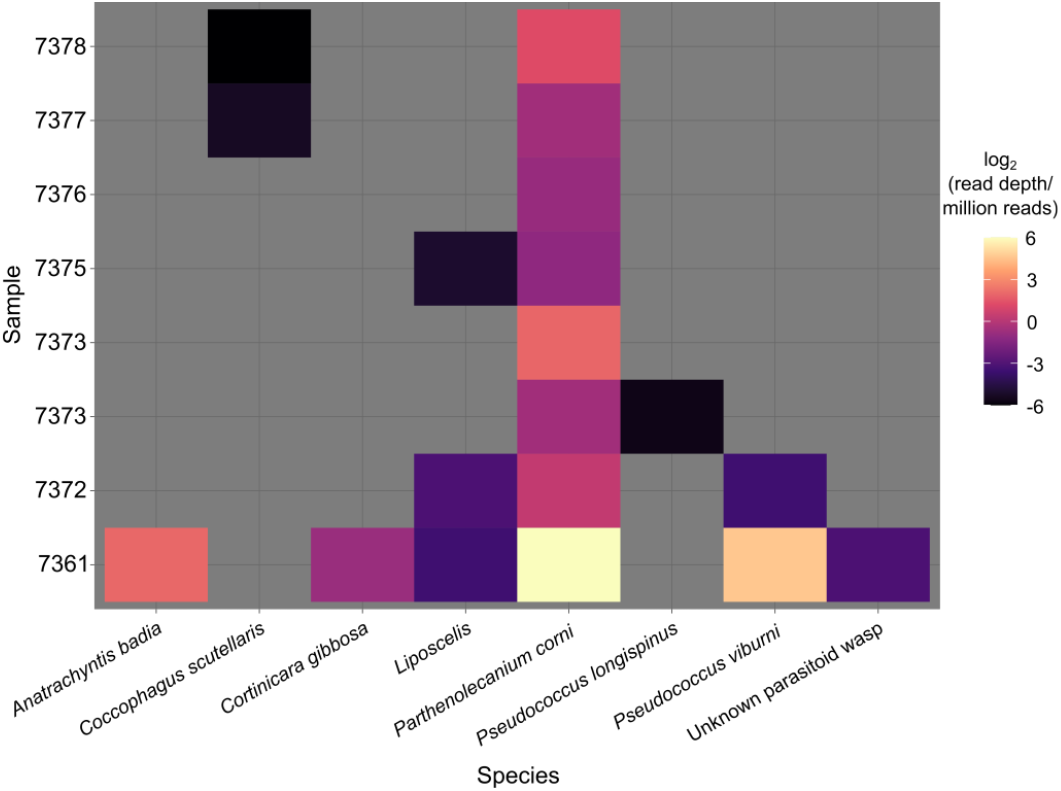
Abundance estimates of insect species present in Australian vineyard samples.

Previous studies have highlighted the presence of several natural enemies of coccids in Australia, including parasitoid wasps, beetles, moth larvae, and lacewings (Rakimov *et al*., 2013; Rakimov *et al*., 2015). Mitochondrial genome fragments were recovered for two distinct Hymenopteran taxa from three different locations. The parasitoid wasp *Coccophagus scutellaris* was identified in the samples from the Barossa Valley, both of which contained *Parth. corni* as the only other insect taxa. A single unknown mitochondrial fragment with best BLAST hits against multiple Chalcidoid wasp mitochondrial genomes was also identified in the 7361 Langhorne Creek sample. The MitoMiner ecosystem monitoring process is therefore capable of determining the abundance of specific scale and mealybug species, while detecting insect species that can act as potential biocontrol agents for these pests. As such, this pipeline could be used as an important biodiversity monitoring tool for assessing the impact of various biocontrol strategies.

Insect abundance estimates based on shotgun sequencing reads mapped against assembled mitochondrial fragments that were classified as insect species using either the BOLD API or by BLASTn against insect NCBI GenBank mitochondrial genomes. Species abundance is presented as the log_2_ of total read depth per million reads in each sample.

In addition to mitochondrial genomes from mealy bug, scale and their potential parasitoids, complete mitochondrial genomes were recovered for the insect species *Anatrachyntis badia* (Florida Pink Scavenger) and *Corticarina gibbosa* (a species of Minute Brown Scavenger Beetle) in sample 7361 from Langhorne Creek (7361), in addition to a partial mitochondrial genome from an unknown species of *Liposcelis* (bark louse) in scrapings taken from the Adelaide Hills (7375), Langhorne Creek (7361) and McLaren Vale (7372). This would suggest these species may also be living within, or associated with colonies of scale and mealybug.

### Evidence for *Parthenolecanium corni* in Australian vineyards

While *Parth. corni* has been described in the Australian taxonomic record (Belbin *et al*., 2021), it has not previously been considered a pest of Australian vineyards. Instead, previous studies into the distribution of soft scale insects in Australian vineyards identified *Parth. persicae* and *Parth. pruinosum* as the only species of *Parthenolecanium* observed (Rakimov *et al*., 2013). Given the ubiquitous detection of *Parth. corni* in the metagenomic samples, additional phylogenetic analysis was carried out on the COX1 sequence against datasets derived from BOLD and NCBI to confirm the MitoMiner classification.

Phylogenetic reconstruction utilizing all available sequences for the *Parthenolecanium* genus across the COX1-5P region, revealed that the *Parth. corni* COX1 7361 haplotype was monophyletic with all other publicly available COX1 sequences for *Parth. corni*, with a maximum genetic distance of 0.9 % across the accessions (Figure 3). Furthermore, the *Parth. corni* 7361 haplotype generated by MitoMiner was identical to sequences obtained from Australian (GBMNA27795-19), Chinese (GBMNC41505-2), and Chilean (GBMIN67815-17, GBMIN67813-17, GBMHH24788-19, and GBMHH24786-19) populations of *Parth. corni* (haplotype H2, Table 2). The two *Parth. persicae* haplotypes present in BOLD were separated by a single base difference, however both had a genetic distance of 9.1 % from the *Parth. corni* 7361 COX1 haplotype (Table 2). In comparison, the single publicly available *Parth. pruinosum* COX1-5P, which was also identical to several sequences from isolates of *Parth. corni*, differed by 0.18 % from the 7361 COX1 haplotype.

**Figure 3:**
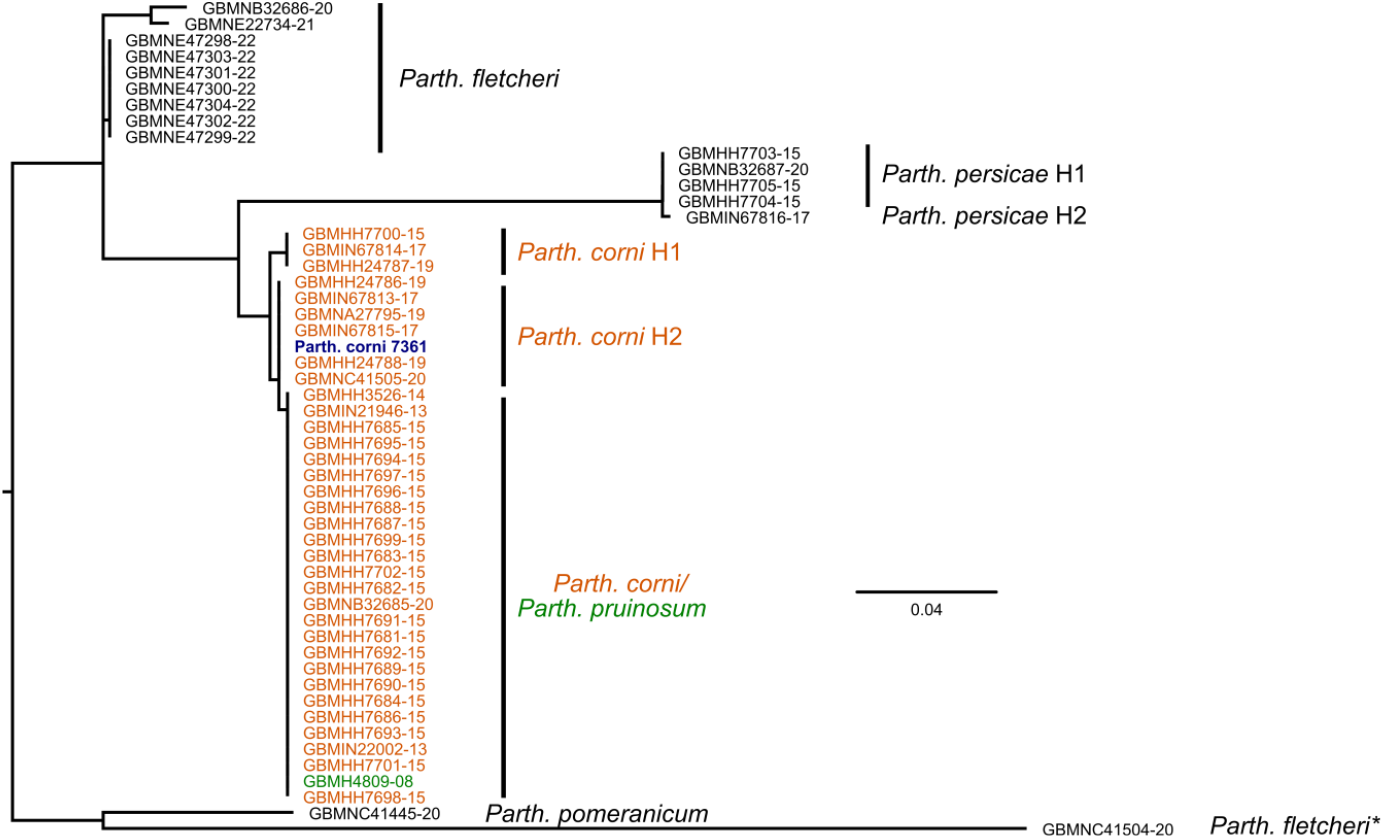
Phylogenetic reconstruction of the COX1-5P locus. Phylogenetic reconstruction of the *Parth. corni* 7361 haplotype against publicly available *Parthenolecanium* datasets across the COX1-5P universal locus sourced from Barcode of Life Database (BOLD). Publicly available data are denoted by their BOLD accessions.

**Table 2.**
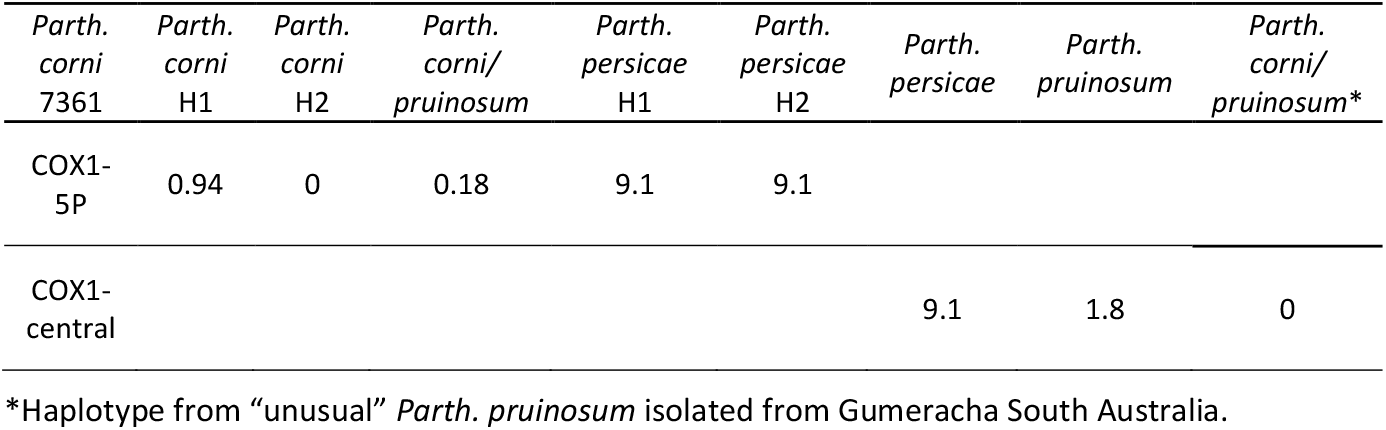
Genetic distance (%) between the *Parthenolecanium* COX1 haplotypes.

While the COX1-5P fragment is the *de facto* standard for molecular-based phylogenetic reconstruction in insects, technical issues necessitated the use of an alternate marker from within the central region of the COX1 mitochondrial gene (COX1-central) in an earlier study by (Rakimov *et al*., 2013). It was therefore not possible for the COX1-central data to be compared against the curated sequences within BOLD (Ratnasingham & Hebert, 2007).

As MitoMiner provided complete mitochondrial genomes for the predicted *Parth. corni* samples in this study, a second phylogeny could be constructed that encompassed sequences form the COX1-central locus produced in Rakimov *et al*. (2013), in addition to a single, independent COX1-central sequence from *Parth. corni* (NCBI:AB439534.1) (Figure 4). As seen for COX1-5P, the *Parth. corni* 7361 COX1 haplotype was highly divergent (9.1 %) from the sequences associated with *Parth. persicae* (Rakimov *et al*., 2013), clearly distinguishing the two species at the molecular level. However, unlike the COX1-5P region, which could not differentiate *Parth. corni* and *Parth. pruinosum*, the 7361 haplotype was shown to be identical to the *Parth. corni* AB439534.1 accession, but distinct (1.8 % difference) from the majority *Parth. pruinosum* samples (Table 2). The exception to this clear species delineation was the presence of sequences from two isolates from Gumeracha, South Australia (KC784922.1 and KC789423.1), that has previously been flagged as representing taxonomically-divergent *Parth. pruinosum* (Rakimov *et al*., 2013).

**Figure 4:**
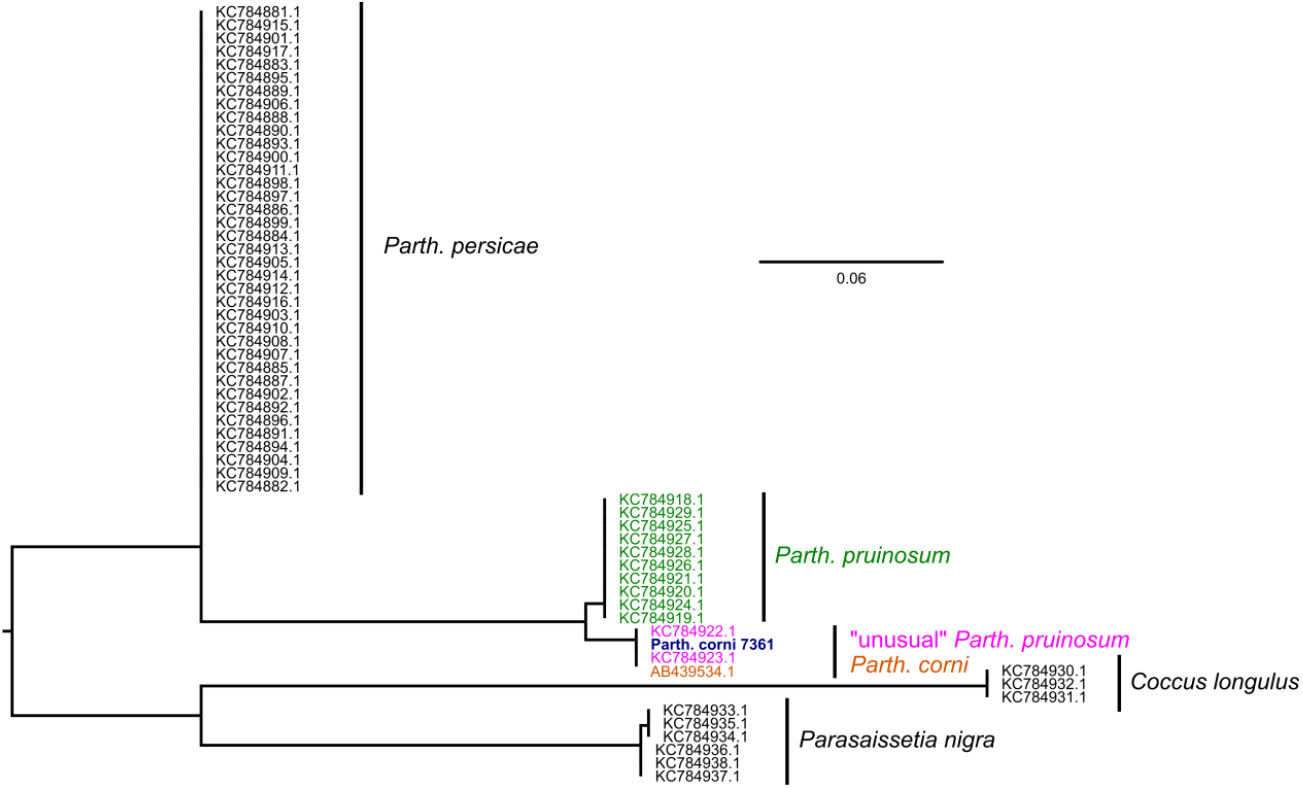
Phylogenetic reconstruction of the COX1-central locus. Phylogenetic reconstruction of the *Parth. corni* 7361 haplotype against publicly available *Parthenolecanium* datasets utilising the COX1-central locus (Rakimov *et al*. 2013). Samples of *Parth. pruinosum* that displayed an unusual taxonomic and lifecycle traits from Rakimov *et al*. (2013) are highlighted in pink. Publicly available data are denoted by their NCBI GenBank accessions.

In addition to providing information from both COX1 loci (5P and central), the availability of a complete mitochondrial genome for the *Parth. sp*. identified in this study allowed for the direct comparison of the two phylogenies. By combining the associated metadata information from both topologies it is likely that the *Parth. corni* species designation provided to the 7361 haplotype by the BOLD classification (which includes vouchered museum specimens and sequences from the Centre for Biodiversity Genomics), is likely to be correct. Furthermore, it also indicates that both Gumeracha sequences are likely to represent *Parth. corni*, rather than “an unusual” representation of *Parth. pruinosum* (Rakimov *et al*., 2013).

Confusion regarding the presence of *Parth. corni* as a major pest within Australian vineyards, is likely due to complications within the *Parthenolecanium* taxonomic record, with several studies questioning the validity of current species limits (Danzig, 1997; Gill, 1988; Nakahara, 1981). Specifically, Gill (1988) stated that *Parth. pruinosum* was indistinguishable in the field from *Parth. corni* if the powdery wax was absent, which is generally the case when dealing with dead and/or dried individuals. However, a lack of tubular ducts on the dorsum, including on the anal cleft, can be used as a taxonomic character to delineate *Parth. corni* and *Parth. pruinosum*. Utilizing these taxonomic characters it was noted that there was a distinct lack of tubular duct along the dorsum of individual soft scale isolated from the South Australian vineyard scrapings, further corroborating that the sequence data obtained from the metagenomic datasets was derived from *Parth. corni*.

The discovery of *Parth. corni* as a major pest within Australian vineyards is consistent with recent vineyard observations of a transition towards a multivoltine mode of reproduction (Venus, 2017). First described in California (Coquillett) *Parth. pruinosum* is generally described as univoltine (García Morales *et al*., 2016; Gill, 1988; Ueda *et al*., 2008), as is *Parth. persicae*, the other major scale species previously reported in Australian vineyards (García Morales *et al*., 2016; Gill, 1988). In contrast, *Parth. corni* is frequently described as multivoltine (García Morales *et al*., 2016), especially in warmer climates (Canard, 1960; Gill, 1988). This has significant implications for the control of scale within Australian viticulture, as current agrochemical applications, which target the scale nymphs, are based around the control of univoltine species (*Parth. persicae* and *Parth. pruinosum*), in which the nymphs present at a single defined time each year. The multivoltine nature of *Parth. corni* dictates that a spread of life stages exist across the year, likely rendering a univoltine-based control strategies far less effective and which could potentially explain the increasing prevalence of serious scale infestations in Australian vineyards.

## Conclusion

Soft scale insects are a major pest in global viticulture due to their ability to facilitate the development of sooty mould and for their propensity as transmission vectors for grapevine viral diseases. The development of the metagenomic-based ecosystem monitoring approach described in this study establishes the ability to detect and quantify major pest species, together with key parasitoids and other associated insect species. The discovery of *Parth. corni* as a significant member of scale infestations provides a key confirmation of empirical observations of scale with multivoltine lifestyles within Australian vineyards and has significant implications for the development of effective control strategies for this important group of pests.

## Acknowledgements

This work was supported by the AWRI and by Australian grapegrowers and winemakers through their investment body Wine Australia, with matching funds from the Australian Government. The AWRI is a member of the Wine Innovation Cluster (WIC) in Adelaide. Special thanks to Marcel Essling for valuable discussions regarding control measures for scale in the Australian Viticultural industry and the grapegrowers that provided access to their vineyards and/or assisted with sample collection.

